# A minimally invasive thrombotic stroke model to study circadian rhythm in awake mice

**DOI:** 10.1101/2024.06.10.598243

**Authors:** Kimberly Marks, Sung-Ji Ahn, Ninamma Rai, Antoine Anfray, Costantino Iadecola, Josef Anrather

## Abstract

Experimental stroke models in rodents are essential for mechanistic studies and therapeutic development. However, these models have several limitations negatively impacting their translational relevance. Here we aimed to develop a minimally invasive thrombotic stroke model through magnetic particle delivery that does not require craniotomy, is amenable to reperfusion therapy, can be combined with in vivo imaging modalities, and can be performed in awake mice. We found that the model results in reproducible cortical infarcts within the middle cerebral artery (MCA) with cytologic and immune changes similar to that observed with more invasive distal MCA occlusion models. Importantly, the injury produced by the model was ameliorated by tissue plasminogen activator (tPA) administration. We also show that MCA occlusion in awake animals results in bigger ischemic lesions independent of day/night cycle. Magnetic particle delivery had no overt effects on physiologic parameters and systemic immune biomarkers. In conclusion, we developed a novel stroke model in mice that fulfills many requirements for modeling human stroke.

## Introduction

Stroke is a leading cause of death and disability worldwide, with a stroke occurring every 40 seconds in the United States alone^1^. Of these, roughly 87% are ischemic, in that brain damage occurs from a clot blocking a major cerebral artery, most commonly the middle cerebral artery (MCA).^2,3^ The standard strategies for reperfusion in acute ischemic stroke include early delivery of recombinant tissue plasminogen activator (tPA) or endovascular thrombectomy, both of which must be performed within hours of a stroke, thus preventing most patients from ever receiving effective treatment.^4^ Furthermore, in roughly 50% of cases, arterial recanalization does not improve functional outcomes (futile reperfusion).^5–9^

Improving upon these existing therapies or developing new ones requires robust stroke models that best recapitulate the human disease. Unfortunately, the various models of MCA occlusion currently available fall short of this goal. The reasons for this include: (a) the production of large infarcts encompassing brain areas outside the MCA territory typically not affected in human strokes, (b) the need for anesthesia, (c) absence of endogenous and/or therapeutic reperfusion, (d) sudden recanalization not reflecting the gradual clot dissolution and reperfusion observed in humans, (e) poor intravital imaging compatibility, and (f) the need for a craniotomy to expose the MCA, which interferes with post-ischemic inflammatory processes and alters CBF dynamics.

Here we describe a novel minimally invasive model of reversible MCA occlusion in mice using intravenous injected thrombin-coated magnetic particles combined with a micro-magnet placed over the intact skull overlying the MCA. We show that this model exhibits several desirable features including applicability to awake or sleeping mice, no requirement for anesthesia during MCA occlusion, slow spontaneous reperfusion akin to that seen in humans, no craniotomy, no unnecessary inflammatory contribution, easy compatibility with 2-photon brain imaging, and reperfusion with tPA.^10–13^ This model is well poised to enable advances in stroke pathobiology, for example the study of circadian effects on stroke outcome and minimizing limitations related the confounding effects of anesthetics and craniotomy, well known translational barriers in preclinical stroke research.^14,15^

## Results

### MCA occlusion by magnetic nanoparticles with endogenous reperfusion

Following a study which used a non-invasive method of reversible MCA occlusions using magnetized nanoparticles^16^, we aimed to develop a stroke model in mice that would reproducibly generate a 85% or greater drop in cerebral blood flow (CBF) necessary to induce ischemic brain injury.^12^ To accomplish this, we placed an attractive magnet over the temporal bone while simultaneously injecting magnetic particles into venous circulation (Fig. 1). A temporary ligation of the ipsilateral common carotid artery (CCA) was necessary to allow for reproducible aggregation of particles at the MCA (Fig. 1a,c). The particles (180nm) used by Jia et al. did not provide the desired 85% or greater decrease in cerebral blood flow to the MCA as shown by Laser Doppler Flowmetry (Fig. 1c). Larger particles (500 nm) provided a drop >85% in cerebral blood flow. However, upon removal of the attractive magnet, reperfusion occurred immediately due to the particles re-entering circulation. This pattern of ischemia/reperfusion is similar to that obtained in the intraluminal filament model, in which reperfusion occurs immediately upon removal of the filament.^12^ To better mimic the gradual endogenous reperfusion observed in patients with ischemic stroke^10,11^, we designed custom magnetic nanoparticles with greater occlusive power. Bovine serum albumin (BSA) coated particles (500 nm) were conjugated to bovine thrombin to form thrombin coated magnetic particles (tMP) (Supplementary Fig. 1a). Eight units of thrombin / mg particles were found to be the limit at which no added thrombin would confer additional potency as determined by a plasma coagulation assay (Supplementary Fig. 1b). Using these tMP, an 85% or greater drop in cerebral blood flow was observed with the added benefit of a more stable occlusion and gradual reperfusion after magnet removal (Fig. 1c, Fig. 2). Longitudinal studies using laser speckle imaging (LSI) revealed reperfusion occurring gradually over several days, matching the perfusion level of the contralateral MCA territory by day 5 after occlusion (Fig. 2a). No CBF reduction was observed in sham animals that underwent the same surgical procedure and tMP injection, without magnet placement apart from the expected roughly 30% CBF decrease from the carotid ligation (Fig. 2b). Infarct volumes 48 hours post-stroke showed a cortical lesion mean±SEM of 12±1.86 mm^3^ (Fig. 1b), while no injury was observed in sham animals. tPA given intravenously 35 minutes after occlusion resulted in recanalization of the occluded MCA, leading to rapid CBF restoration to near baseline levels and in significantly smaller injury volumes (2.7±2.2 mm^3^) (Fig. 1d,e). No significant difference in infarct volumes was observed when comparing male and female mice (Supplementary Fig. 2).

**Figure 1.**
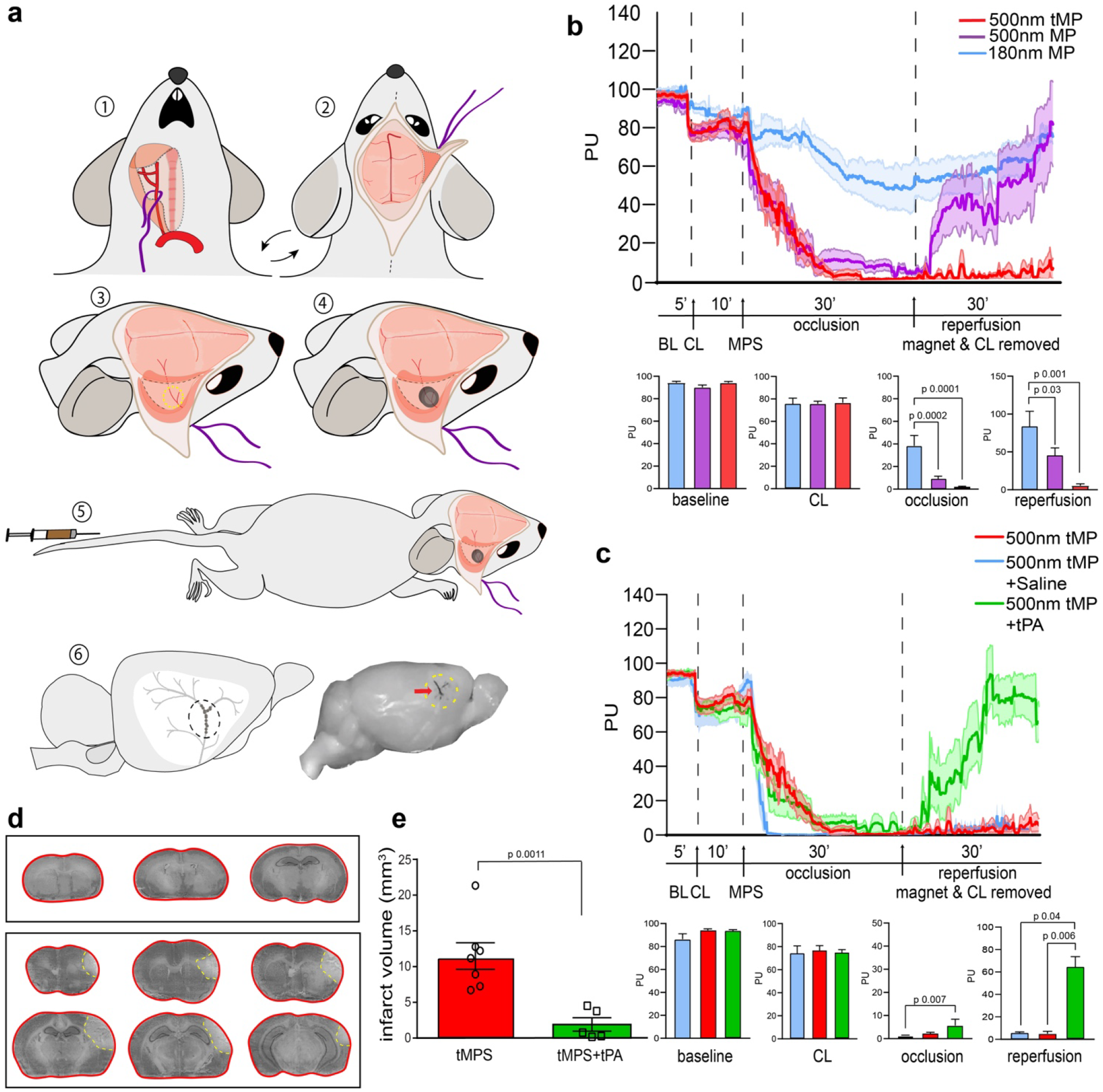
Surgical scheme and assessment of magnetic particle stroke. **a,** Surgical strategy for MCA stroke using magnetic particles: 1. Right common carotid artery surgically isolated and held by nylon thread; 2. Animal turned supine and scalp excised to remove right temporal muscle, exposing temporal bone; 3. Thinning of temporal bone to ∼70 µm over MCA using burr drill; 4. Placement of magnet over thinned area; 5. Tail vein injection of 250 µl tMP in saline; 6. Accumulation of particles to MCA. Upon removal of magnet, particles appear discretely in the distal branches of the MCA. **b,** Laser doppler flowmetry measuring relative cerebral blood flow (CBF) changes as perfusion units (PU) in the MCA territory before surgery, following carotid ligation (CL), tMP injection (MP), 30 minute occlusion and reperfusion after the removal of both magnet and CL. Effect of particle size and composition on LDF (180nm n=5, 500nm MP n=9, 500nm tMP n=9; bar graphs show n=5-9 binned time points per group. Occlusion and reperfusion groups showing binned data points from final 10 minutes of recordings). 180nm particles failed to provide sufficient decrease in CBF. 500nm particles provided sufficient drop in CBF with immediate reperfusion upon magnet removal. 500nm thrombin particles (tMP) provide desired sustained drop in CBF following magnet removal. Data are mean±SEM, Kruskal-Wallis test. **c,** LDF trace of 500nm tMP with and without administration of bolus 10mg/kg tPA modeling therapeutic reperfusion. (500nm tMPS alone n=9, 500nm tMP with tPA n=5; bar graphs show n-5-9 binned time points per group. Occlusion and reperfusion groups showing binned data points from final 10 minutes of recordings) Sham group performed as 500nm tMP with administration of bolus of physiological saline (n=4). Kruskal-Wallis test performed on binned-segmented CBF (final 10 minutes). Note all groups resulted in expected 85% or greater drop in CBF. **d,** Cresyl Violet staining of 30 µm thick coronal brain sections collected at 300 µm intervals. Infarct outlined in yellow. Top panel is sham (magnet and carotid ligation surgery without particles). **e,** Infarct volume 48 hours with and without 10mg/kg tissue plasminogen activator (tPA). Individual values and mean±SEM are shown. Unpaired t-test (n=5-7/group).

**Figure 2.**
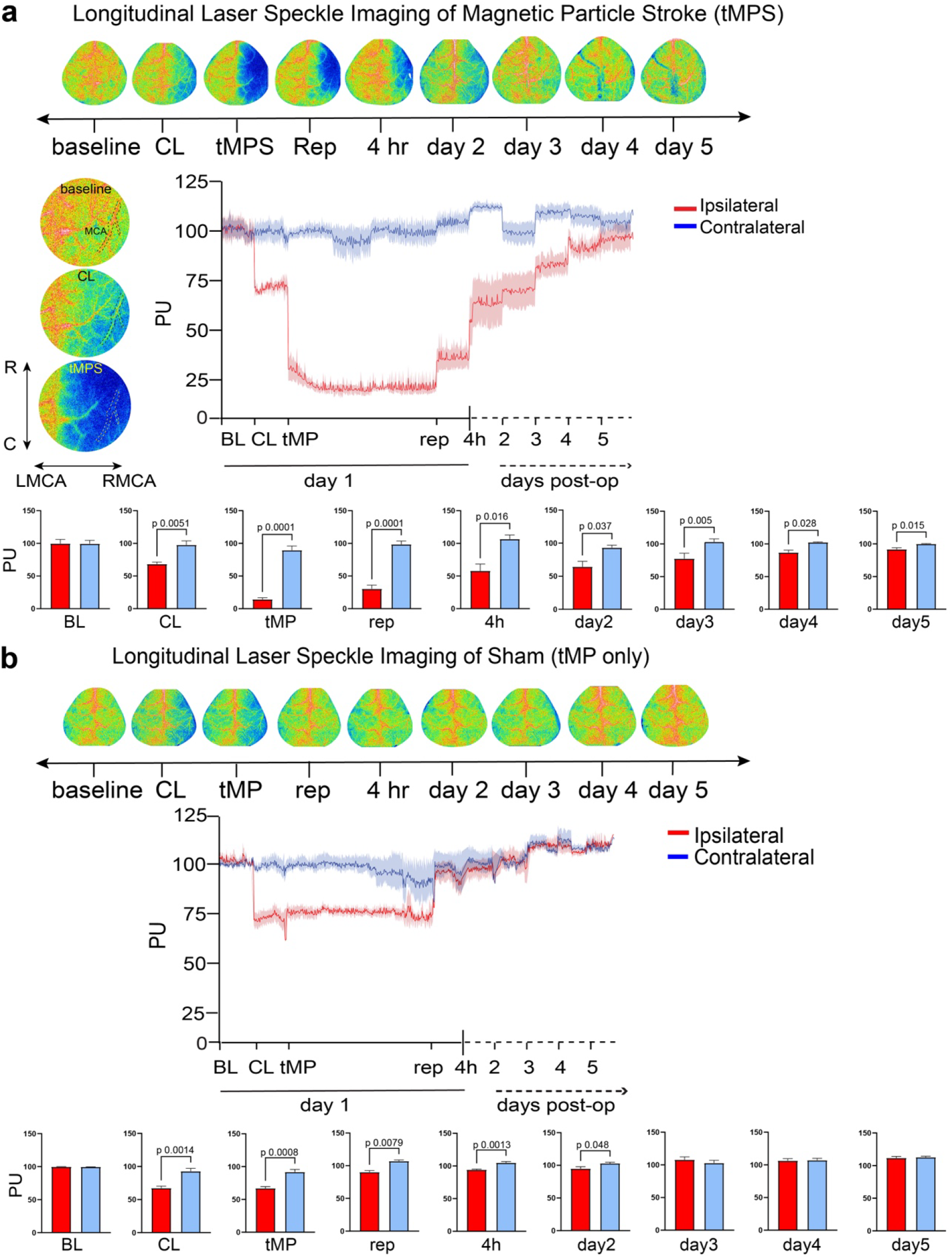
Longitudinal imaging of tMP stroke and reperfusion. **a,** Longitudinal Laser Speckle Imaging during surgery and five days following stroke. Cerebral Blood Flow (CBF) as perfusion units (PU) of ipsilateral and contralateral middle cerebral artery/affected cortex: Baseline (BL) 10 minutes, CCA ligation (CL) 10 minutes, tail vein administration of 500nm thrombin magnetic particles (tMP) 30 minutes occlusion. Magnet and carotid ligation removed (Reperfusion) 10 minutes. Values were normalized and binned for last 5 minutes for all stages with exception of final 10 minutes for tMP injection. tMPS shows a gradual increase of CBF to be restored to ipsilateral cortex by day 5; n=6/group, unpaired-t test. Values are mean±SEM. **b,** Sham surgery (tMP but no magnet) shows CBF (PU) restored after CL is removed; n=5/group, unpaired-t test. Values are mean±SEM. Left of top trace shows zoomed-in representative images with right middle cerebral artery (RMCA) outlined by dashed line. R, rostral; C, caudal; LMCA, left middle cerebral artery.

### Particle body distribution, impact on blood chemistry and inflammatory markers

Next, we used body and brain T2-weighted MRI imaging of sham mice injected with tMP or saline and tMPS (stroke) mice to determine the fate of the particles. Particles were found largely in the liver 1 hour post injection (Supplementary Fig. 1c). Hyperechoic intensity in the brain resulting from accumulation of magnetic particles to the MCA and areas of cortical injury were captured by coronal MRI (Supplementary Fig. 1d). To determine the impact of the particles on liver function, we assessed serum alkaline phosphatase, alanine transaminase, aspartate aminotransferase, total protein, triglyceride, and total bilirubin at 48 hours, 7 days and 14 days following tMPS and in sham-treated mice (Supplementary Fig. 3a). No statistically significant differences were found in groups between days except for ALP showing a moderate decrease in sham tMP and tMPS groups. Arterial pH, pCO_2_, pO_2_, and mean arterial pressure measured before and immediately after tMPS and tMP sham surgery showed no statistical difference between groups (Supplementary Fig. 3b). pH, pCO_2_, pO_2_ were all within normal physiologic ranges for tMPS and tMP sham.^17^

To rule out systemic inflammation resulting from tMP, selected serum cytokines known to be regulated in ischemic stroke were measured at 4, 24, 48 hours, and 7 days following tMPS, tMP sham surgery, or saline-treatment.^18–21^ As expected, given the inflammatory sequelae of stroke, serum cytokines were increased in tMPS when compared to control groups (Supplementary Fig. 3c).^18,19,22^ No differences were found between tMP sham and saline sham groups. We conclude that the tMP themselves do not elicit a systemic inflammatory response that could aberrantly contribute to ischemic injury and recovery and that any significant impact on systemic inflammatory markers results from the ischemic injury itself.

### Behavioral deficits after tMP stroke

Given that the somatosensory cortex is the major brain area targeted by distal MCA occlusion, we performed neurobehavioral tests that predominantly test the function of this brain region.^23^ Wire hanging measures grip strength, balance, and endurance with the behavioral output as latency to fall from the suspended wire.^24^ Mice receiving tMPS had shorter times suspended when compared to tMP sham groups, a deficit that could be rescued by therapeutic doses of tPA (Fig. 3a). The corner test detects sensory and motor asymmetries related to barrel cortex outputs, limb coordination and postural biases.^25^ Mice receiving tMPS had an 80% preference to turning right, ipsilateral to the side of the infarct. These findings were statistically significant compared to tMP sham animals and could also be rescued using tPA (Fig. 3b). The adhesive removal test is used to assess cortical damage and coordination deficits.^26^ At 3 days following injury, mice receiving tMPS had increased latency to contact the adhesive attached to the paw and increased time to remove adhesive when compared to the tMP sham group (Fig. 3c).

**Figure 3.**
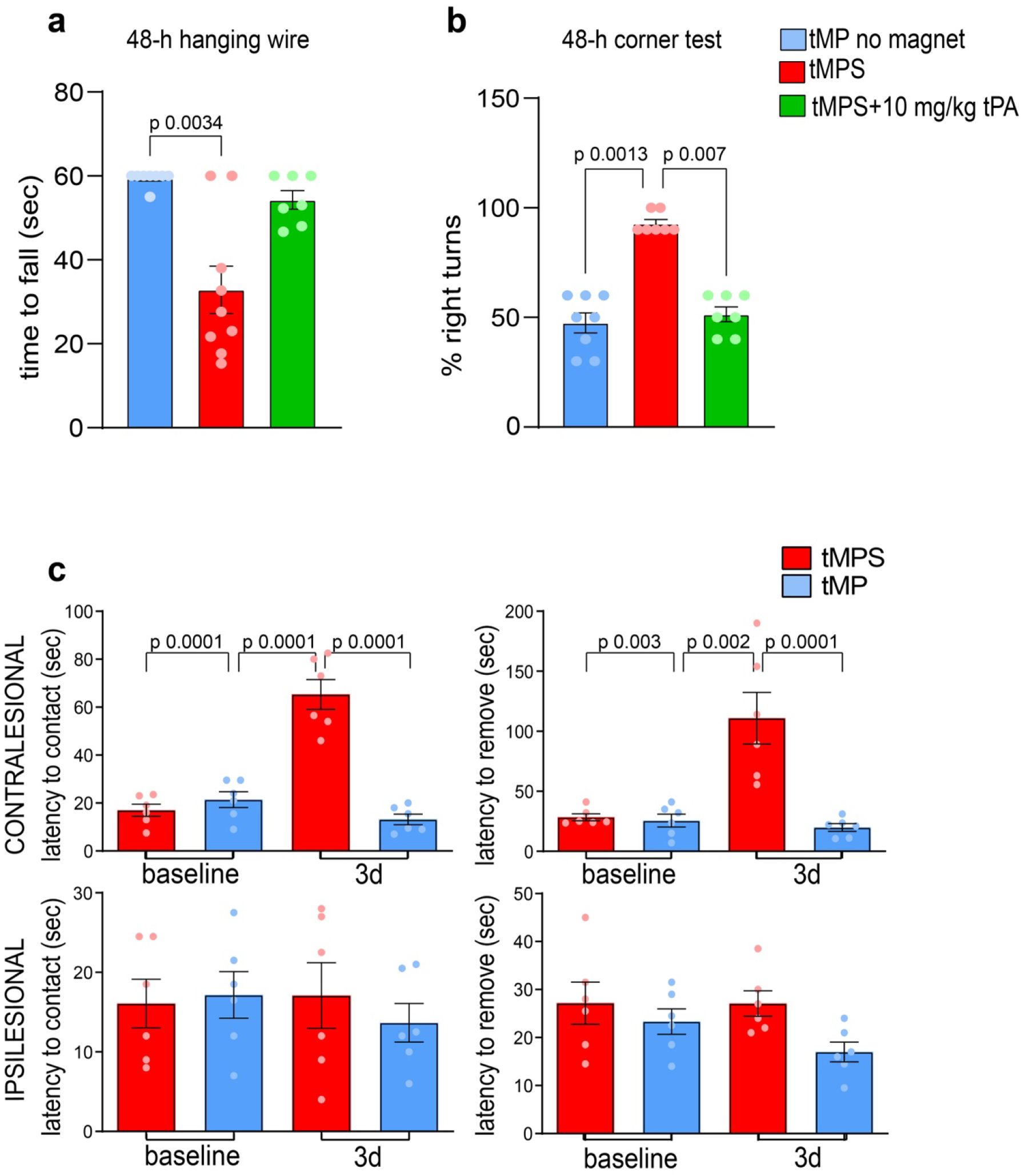
Motor-sensory impairment following thrombin magnetic particle stroke (tMPS). **a,** Hanging wire test shows motor, grip strength and balance impairments in MP stroke compared to sham mice at 48 hours. Kruskal-Wallis test. **b,** Corner tests shows sensorimotor perception impairments in MP stroke compared to sham at 48 hours. Kruskal-Wallis test. **c,** Time-to-contact and time-to-removal adhesive tape test between sham (receiving particles but no magnet) and stroke mice (particles and magnet). Notice significant increased time to remove adhesive at 3 days following stroke. One-way ANOVA with Tukey’s multiple comparisons test. Individual values and mean±SEM are shown.

### tMP stroke results in immune cell infiltration in the neocortex

Neuronal cell loss, glial activation, and immune responses of brain-resident and peripheral immune cells are hallmarks of stroke pathology.^22,27,28^ To visualize cytologic changes induced by tMPS, we performed immunocytochemistry 48 hours post stroke showing axonal, microglial, and astrocytic changes (Supplementary Fig. 4a). MAP-2 staining showed axon depletion in the infarct core, Iba-1 staining showed a reduction of microglia/macrophages in the core, while GFAP-positive astrocytes were increased in the peri-infarct region at 48 hours. Staining of the distal MCA branches using fibrinogen and platelet marker CD41 showed the presence of platelets and fibrinogen in distal branches of MCA four hours after stroke, with labeling of penetrating vessels in the parenchyma (Supplementary Fig. 4b).

We then used flow cytometry to examine in greater detail the immune cell populations at 1, 2, and 7 days after tMPS and comparing to the uninjured contralateral cortex.^29,30^ (Fig. 4). CD45^hi^ infiltrating leukocytes were elevated in the ipsilateral cortex at all time points. No significant changes in CD45^int^ (microglia) and endothelial cells were found over time. Ly6C^hi^ and Ly6C^lo^ monocyte-derived cells (MdC) were elevated at 48 hours and Ly6C^lo^ MdC at 7 days. Neutrophils were elevated at 48 hours in the ipsilateral cortex and T cells were most elevated at 7 days when compared to the ipsilateral cortex at 1 day and the contralateral cortex at 7 days. Finally, NK cells were elevated at 7 days. B cells did not differ between groups, but levels were elevated in the ipsilateral cortex (Fig. 4).

**Figure 4.**
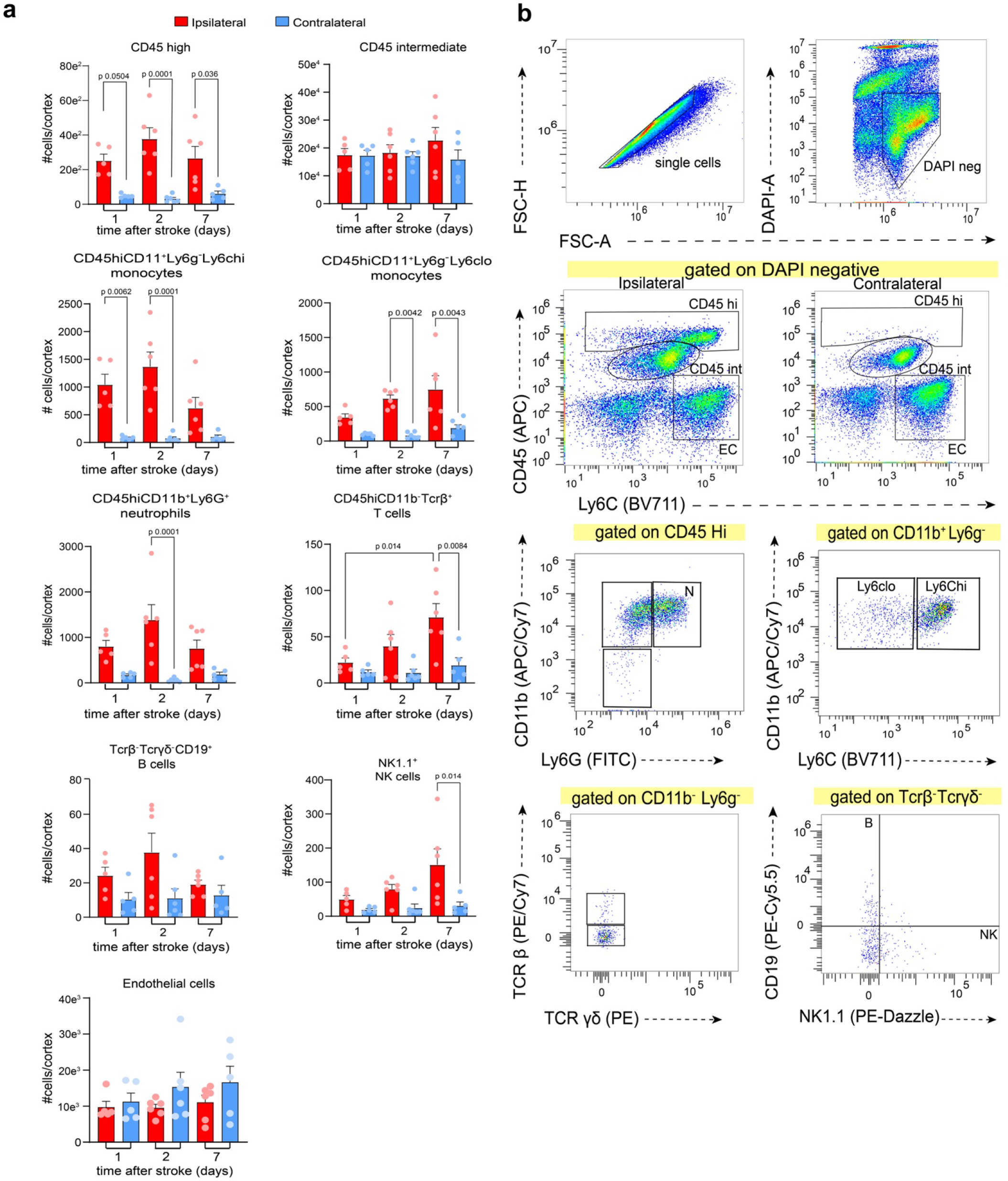
Immune response to tMP stroke. Following stroke, animals were perfused transcardially using 1X PBS followed by an enzymatic digestion of cortical tissue for FACS sorting with antibodies: CD45 (APC), Ly6C (BV711), TCRγδ (PE), NK1.1 (PE-Dazzle), CD19 (PE-Cy5.5), TCRβ (PE-Cy7), Ly6G (FITC), CD11b (APC/Cy7), DAPI**. a,** Cell counts per cortex at 1, 2, and 7 days following stroke (n=6/group) One way ANOVA. **b,** Gating strategy flow scheme is shown for cortices at 2 days. Plots gated on DAPI^−^ and CD45^hi^ compare both hemispheres. Subsequent plots below show ipsilateral cortex only. EC, endothelial cells; N, neutrophils; T, T cells; B, B cells; NK, natural killer cells.

### Intravital imaging of tMP stroke

The tMPS procedure can be modified to accommodate a non-invasive thin-skull cranial window and headplate for multiphoton imaging.^31–33^ Intravascular flow was assessed by measuring red blood cell velocity using line scanning at baseline, following carotid artery ligation, injection of tMP, removal of magnet and ligation (Supplementary Fig. 5a). Red blood cell velocity slowed down following tMP administration, with the injected particles seen as white specs (Supplementary Fig. 5c). By 20 minutes and following magnet removal, movement of red blood cells is completely stalled (Supplementary Fig. 5b,c). Diameter measurement of MCA distal branches was decreased during tMPS (Supplementary Fig. 5d). The movies obtained at baseline and following tMP injection (Supplementary Movies 1-5) show the vasodynamic changes MCA branches downstream of the magnet. Frequent oscillations of constriction and dilation, blood turbulence, and reversal of blood flow were seen in response to tMPS.

### tMPS in awake mice

Most models for MCA ischemia/reperfusion *in vivo* require anesthesia, which is not the case for human stroke.^34,35^ Anesthetic agents have a profound impact on the brain and are endowed with neuroprotective effects.^14,15^ To eliminate the confounding effect of anesthesia on the development of ischemic brain injury, the tMPS model was adapted to induce strokes in the awake state. To eliminate the need for transient CCA, a micro-coil was placed around the ipsilateral CCA to induce the CBF reduction needed for the microparticles to aggregate and occlude the MCA in the presence of the magnet. This technique also connects the tMP model to carotid artery disease and stenosis, which moreover, has a sizable contribution in the probability of developing an ischemic stroke.^36,37^ Animals were trained to tolerate awake tail vein restraint for injection of tMP in days leading up to stroke, minimizing the effects of stress on stroke pathophysiology. Animals were briefly anesthetized for coil and magnet placement before being allowed to recover for awake injection of particles (Fig. 5a). This roughly 25-minute surgical procedure is not thought to provide animals with anesthetic preconditioning that would impact development of the infarct.^14,38^ Compared to tMPS with temporary carotid ligation (method described above), tMPS with unilateral carotid stenosis (designated tMPS coil) resulted in similar infarct volumes and behavioral outcomes (Fig. 5b). Longitudinal LSI imaging of the mice undergoing tMPS coil procedure showed similar reperfusion dynamics when compared to the tMPS animals (Fig. 5d).

**Figure 5.**
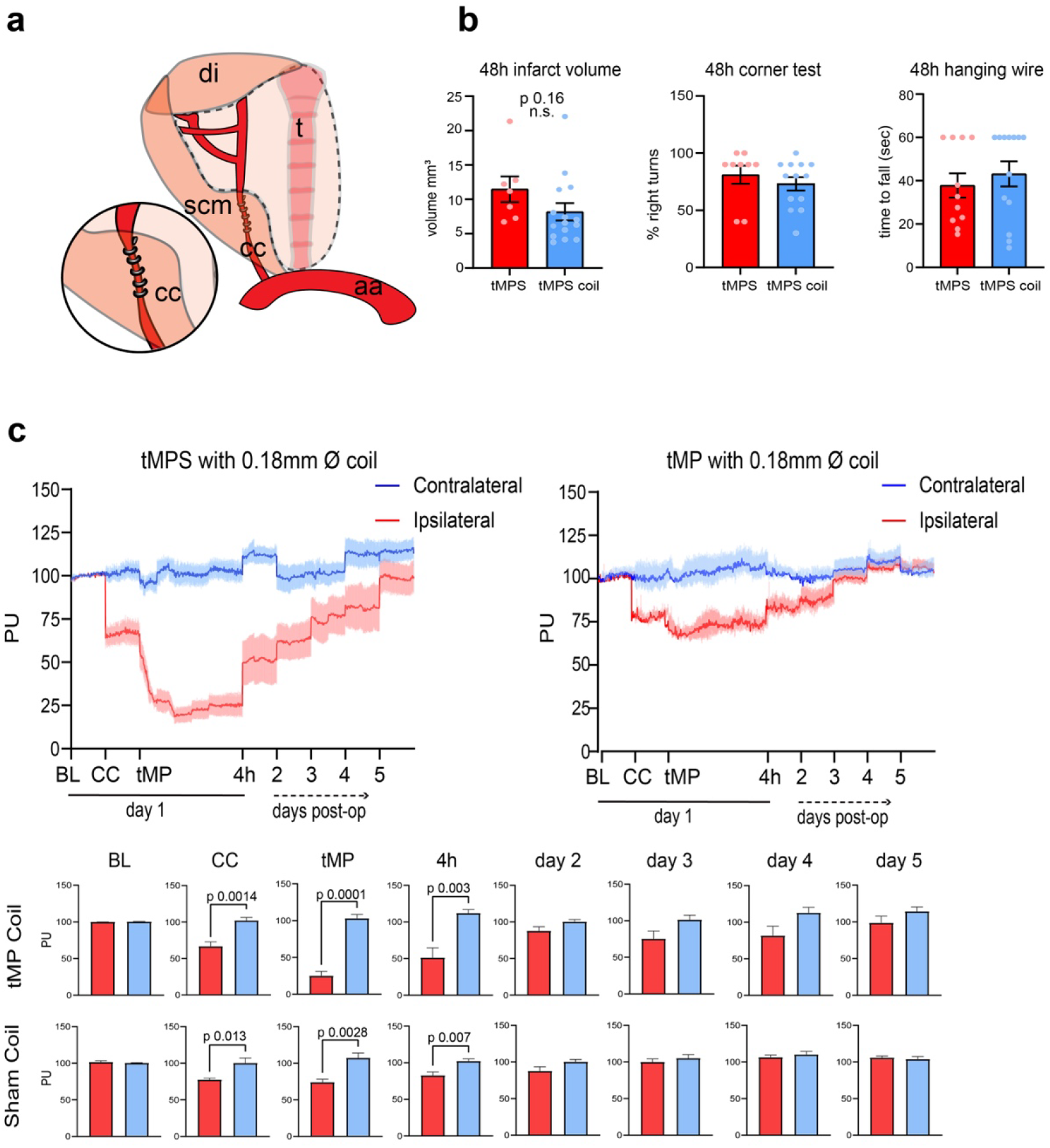
tMP stroke (tMPS) model using permanent unilateral coil and magnet. Animals underwent a short surgery (∼25 minutes) under isoflurane (1.5%) for permanent placement of carotid coil and magnet (as outlined in methods) followed by injection of thrombin particles. **a,** Instead of temporary ligation with nylon suture, animals received unilateral permanent placement of 0.18mm Ø coil to the carotid artery that is ipsilateral to cortex of intended injury and magnet placement (magnet left in place). di, digastric muscle; scm, sternocleidomastoid muscle; aa, aortic arch; cc, common carotid artery; t, trachea. **b,** Infarct volume of tMPS compared to tMPS using coil both under anesthesia, unpaired t-test, p-value. 48-hour corner and hanging wire behavior test between awake tMPS and anesthetized tMPS were not significant, unpaired t-test. **c,** Animals were anesthetized (1.5% isoflurane) for surgery and LSI measurement. Animals were monitored for CBF as perfusion units (PU) following a baseline reading (BL), the unilateral coil placement on the carotid ipsilateral of permanent magnet placement (CC), following injection of particles and consequent tMP stroke (tMPS), 4 hours following stroke and up to 5 days. Sham animals followed identical surgical procedure and particle injection except for magnet placement (tMP). n=5/group, unpaired t-test.

Next, we compared infarct volumes produced by tMPS coil with anesthesia to tMPS coil in awake animals. We found that infarct volumes were significantly smaller with anesthesia than without anesthesia (Fig. 6a). Since clinical and experimental data indicate circadian variations in the magnitude of stroke injury and treatment, we induced stroke in day-time and night time.^39–41^ We found no significant infarct volume difference between day-time and night-time strokes in either anaesthetized or awake animals (Fig. 6a). Finally, surgery-to-injection time intervals showed no correlation between the groups receiving surgery first versus last and overall stroke volume (Fig 6b,c). This, along with the longitudinal LSI data (Fig 5c), indicates that the carotid stenosis and CBF reduction by micro-coil provides a stable drop in CBF within this time frame of 2-3 hours from surgery to injection for the particles to aggregate appropriately.

**Figure 6.**
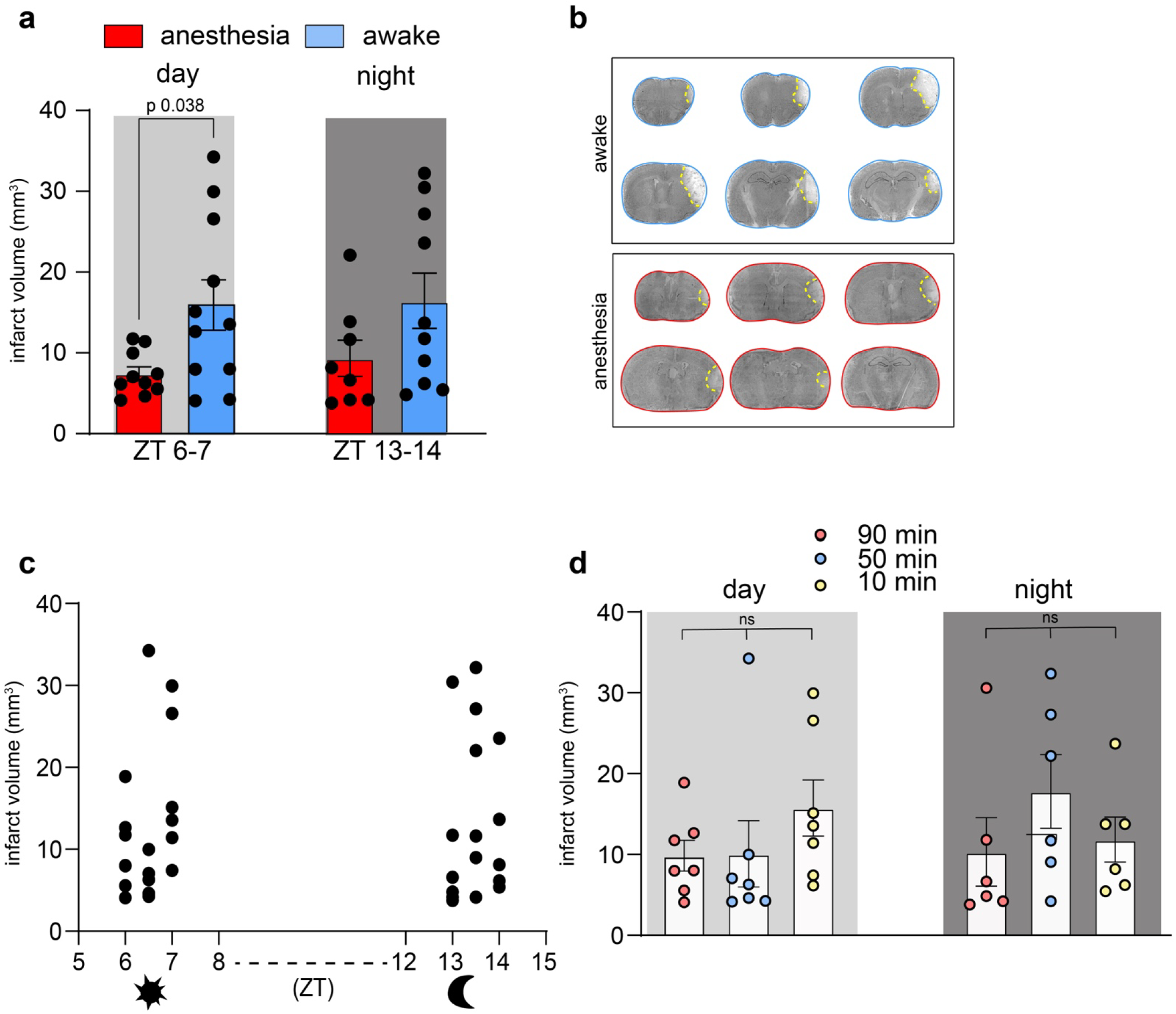
Effects of anesthesia and circadian cycle on tMP stroke. a, Coil and magnet placement surgeries were performed mid-morning and injected as one group following zeitgeber time (ZT) 6-7 or in evening at (ZT) 13-14. Unpaired t-test. **b,** Cresyl violet staining shown of 48hr post awake daytime coil tMPS, (top) outlined in blue and 48hr post isoflurane anesthetized daytime coil tMPS (bottom) outlined in red. Infarct territory outlined in yellow. **c,** Paired time-to-particle injection following coil and magnet surgery. Animals were injected in the order of surgery completed within 1-2 hours of full recovery. **d,** Infarct volumes of mice shown as time from end of surgery to particle injection. Individual values and mean±SEM are shown. One-way ANOVA with multiple comparisons shows no effect of time between surgery and stroke induction.

## Discussion

Despite recent progress in therapeutic interventions, ischemic stroke remains a major risk factor for death and long-term disability. While preclinical stroke models have been pivotal in elucidating pathomechanisms and identifying potential druggable targets, the translation of these findings into clinical therapies has largely failed. The majority of preclinical stroke studies in mice have been performed by transient intraluminal proximal MCA occlusion (tMCAo) which blocks blood flow to the entire MCA territory and by permanent distal MCA occlusion (pMCAo) specifically targeting cortical MCA supply territories. While tMCAo can mimic reperfusion and is therefore preferred when modelling large vessel ischemic strokes in humans that have been recanalized by tPA treatment or mechanical thrombectomy, it also reduces blood flow to the lenticulostriatal arteries leading to ischemic injury of basal ganglia, a condition rarely seen in large vessel ischemic strokes in humans.^35,42^ The pMCAo model results in lesions restricted to the neocortex and is therefore a better mimic of ischemic MCA strokes in humans but can’t be reperfused by either pharmacologic or endogenous thrombolysis.

To address these limitations, several focal thrombotic and thromboembolic stroke models have been developed and performed in mice. These models include photothrombotic (PT) stroke, topical FeCl_3_ application to the MCA, and intraluminal thrombin injections. Similar to the tMPS model, the PT stroke model is minimal invasive but does not mimic large vessel occlusion and is characterized by endothelial injury and vasogenic edema. Moreover, the mechanism of thrombosis in this model are distinct from classical thrombus formation as they do not require the presence of platelets or activation of the classical coagulation cascade.^43^ Local application of FeCl_3_ over the MCA requires craniotomy and produces a platelet-rich thrombus that is not responsive to tPA reperfusion.^44^ Intraluminal injection of thrombin into the MCA instead generates a thrombus that is composed of platelets and fibrinogen and, similar to the tMPS model, is amendable to tPA thrombolysis.^45,46^ The surgical procedure, however, requires craniotomy and the grade of clot formation is not consistent.

More recently, stroke models based on delivery of synthetic particles to the MCA or brain have been developed. Untargeted delivery of microspheres or cholesterol crystals via the CCA produce permanent microvessel occlusion resulting in randomly distributed microinfarcts in cortical and subcortical brain regions that do not resemble large vessel strokes.^41,47,48^ In contrast, magnetic nanoparticles can be directed to a determined vascular target by attractive magnet placement. One such study used 100nm thrombin-linked magnetic particles attracted to a NdFeB magnet placed on the CAA.^49^ Upon clot formation and removal of the magnet, the thrombus embolizes to the MCA leading to vessel occlusion. To date, no data are available detailing the extend of the ischemic territory, infarct volumes, reproducibility, or the response to rtPA. A second study made use of magnet placement on the MCA paired with magnetized red blood cells in P0-7 pups modeling perinatal cerebral hemorrhage and stroke.^50^ Most recently, another group used magnetic particles to induce infarcts to the MCA P6 pups and P28 mice.^51^ While these studies show promise and confirmation of what we see to be powerful tool using nanoparticles for modeling cerebral ischemia, neither is modeling ischemic stroke in awake mice. Additional comparisons made between experimental models are shown in Supplementary Table. 2.

The tMPS model of stroke has several advantages over existing models. First, and most compelling, this model allows for stroke to occur in awake, non-anesthetized mice. The initial method development made use of anesthesia to validate and compare against the existing metrics of cerebral blood flow, behavior, and the inflammatory response of the established models which also use isoflurane. However, the model has been adapted to introduce MCA occlusion while the animal is fully conscious.

Modeling MCA stroke in non-anesthetized mice has long been a goal of the field and would eliminate the potentially confounding neuroprotective impact of inhalant anesthetics.^14,15,52,53^ This awake model offers other technical benefits to experimenters: for example, it allows investigators to conduct strokes during the animal’s wake cycle, having prepared the animals during the surgeons’ daytime work hours and inject with tMP early evening at the start of the mouse’s wake cycle without needing to perform stroke surgeries at nighttime. This should allow to model morning-onset strokes frequently seen in humans.^39,40,54–56^ Placing a coil around the carotid and a magnet over a thinned temporal bone requires roughly 20 minutes of surgery time, with the injection itself taking no more than 1-2 minutes to accomplish and several animals can be injected with tMP all at once, as a batch. In contrast to surgery under anesthesia that requires an average of 50-60 minutes minimum to complete (namely due to requiring surgeons to wait 30 minutes following injection of the particles before removing the magnet and reversing the ligation), preparing animals for awake strokes ultimately saves time.

A major finding of this study is that infarct volumes were significantly smaller in tMPS animals with micro-coils injected under anesthesia compared to those injected awake, possibly indicating protective effects of anesthesia. No differences were seen in the infarct volumes or outcomes of these two groups when comparing strokes performed during the active or inactive cycle.

Animals under anesthesia undergoing tMPS either by temporary carotid ligation or by permanent coil placement resulted in similar infarct volumes and behavioral outcomes (Fig 5a,b). Interesting to note is that following the permanent placement of a micro-coil, the magnet and after particle injection, reperfusion to the affected MCA closely matched what was seen with the reversible ligation and magnet removal in anesthetized mice, indicating that for an infarct to occur the magnet and carotid stenosis either by coil or ligation does not need to be removed after particle injection and that reperfusion occurs regardless of their removal (Fig. 5d).

This endogenous reperfusion is another marked advantage of the magnetic stroke model. Spontaneous recanalization and vascular patency after several days is a characterized phenomenon that occurs in stroke patients that do not undergo rtPA treatment.^11,42^ Additionally, human ischemic stroke often has partial spontaneous reperfusion after endogenous dissolution of the thrombus within the first 48 hours following stroke.^10,11^ In a study following patients with MCA strokes by transcranial doppler within 6 hours of infarct and at 1 week, more than half of the patients had normal flow to the MCA by 7 days with 3 to 9 month follow-ups showing the same findings.^57^ This is similar to the reperfusion pattern being shown by this model.

Large vessel atherosclerosis and carotid stenosis are the largest contributors to the pathophysiology of thromboembolic stroke to the MCA, accounting for most stroke cases in humans.^36,37,58^ Logically the primary cause of injury is relevant to the treatment of acute ischemic stroke in humans, thus animal models should also be able to address this issue. The magnetic particle model models thrombotic stroke with the injection of occlusive particles into mouse circulation, allowing for aggregation of thrombin particles in a clot-like fashion to the MCA below the attractive magnet, it also has the capacity to model carotid stenosis using a unilateral carotid micro-coil.

Our studies following injection of tMP measuring blood gases, liver enzymes and cytokines showed minimal impact to overall physiological parameters in mice. Iron (II,III) oxide particles have been used as imaging agents in humans providing good biocompatibility and negligible toxicity.^59^ Dextran and iron oxide magnetic nanoparticles in particular have been used for diagnostic imaging and therapeutics for cancer in humans.^60^ ^61^

Given the increased interest in the roles of meninges regulating brain function and the inflammatory response after stroke^62–64^, the tMPS model, not needing craniotomy and compatible with in vivo imaging, is ideally suited to address their roles in the response to thrombotic stroke.

Finally, limitations to this model are few. The largest limitation of the tMP stroke model is technical. While commonly done, injection by tail vein can be a challenge for new surgeons. One should be very proficient in injecting by tail vein (particularly in awake mice) before starting to perform the technique.

In summary, here we present a model of thrombotic stroke of the distal MCA territory leading to reproducible cortical infarcts that does not require craniotomy, can be performed in awake mice, is amendable to rtPA reperfusion, and shows neurobehavioral and immune response characteristics similar to the ones reported for more invasive distal MCA occlusion models. We hope that the tMP stroke model will close some gaps in modeling human stroke in the preclinical setting.

## Materials and Methods

### Animals

8-12 week old C57Bl/6J male mice from JAX laboratories were used for all studies. Where indicated, 19-24 week old C57Bl/6J female mice were used. Mice were housed socially (three to five mice per cage) in individually ventilated cages, with *ad libitum* access to food and water and under controlled conditions (22 ± 2 °C, 12:12 h light/dark cycle with light phase from 7:00 a.m. to 7:00 p.m.; 40–60% humidity). Mice from the same cage were subjected to stroke or sham surgeries alternating respectively and returned for recovery. Post-surgery sham and stroke mice were housed together. Surgical success and attrition are presented in Supplementary Table 1.

### Magnetic particle size and composition

We used magnetic particles that are a composite of magnetite [iron(II,III) oxide] and 40kDa dextran matrix (Nanomag-D, 180nm or 500nm, 10 mg/ml, MicroMod, Germany Cat No. 09-82-182 and 09-00-502). When paired a Neobydium attractive magnet (1.59 x 0.79mm thick N52 NdFeB axially magnetized Ni coated magnet, Zigmyster magnets Cat No. D116-132N52) placed on the temporal bone, particles accumulated to the MCA after intravenous tail vein injection. Particles were rinsed with a magnetic pull-down rack three times using physiological saline and resuspended in 250 µl to make 10mg/ml. Particles were sonicated in a water bath for 10 minutes immediately prior to injection.

### Thrombin composite particles (tMP)

Custom order nanoparticles (MicroMod Nanomag-D, albumin (BSA), 500nm, 10mg/ml) were conjugated to bovine thrombin (Millipore) to induce clot formation. 20 U bovine thrombin (Millipore Sigma Cat No. 605157) in 0.1% Bovine Serum Albumin (BSA) was added directly to aliquoted doses of 250 µl suspended magnetic particles and incubated overnight at 37°C. Following incubation and using a magnetic bead separation rack, supernatant was discarded, and particles were rinsed twice with using 300 µl saline and magnetic pull down again to discard supernatant. Thrombin nanoparticles were resuspended in 250 µl of physiological saline and sonicated in bath sonicator for 10 minutes before injection. The dose of thrombin conjugated to nanoparticles was determined by a coagulation assay. 5 U thrombin increments were added to microcentrifuge tubes containing 50 µl of human citrated plasma (Sigma Cat No. P9523-1ML), the time to clot formation was recorded. The same 5 unit increments of thrombin were added to magnetic nanoparticles (250 µl) and incubated overnight at 37°C. After washing the particles, human citrated plasma (Sigma Cat No. P9523-1ML) was added and time to clot formation was recorded.

### Magnetic particle model of stroke

Mice were anaesthetized with isoflurane (4.5-5% induction, 1.5% maintenance). Bupivacaine 0.25% solution was administered trans-dermally prior to any incision at both scalp and neck. Toe-pinch reflex was done to check for anesthetic depth. Body temperature was maintained at 37°C using a feedback rectal probe and heating pad.

#### Reversible CCA ligation

With animal supine, a 1cm incision was done to shaved and sterilized neck skin over trachea, without cutting trachea or muscles. Using sterile cotton swab soaked with saline, neck muscles were gently separated to reveal trachea and surrounding vessels. The right CCA was isolated using sterile micro-forceps. Care was taken not to disturb the vagus nerve, which can be seen as a fine white strand behind the carotid. Once the vessel was isolated, a 10cm long 6-0 US silk surgical suture was fed below the carotid. Both ends of the suture were left loose and open until ready to inject particles. Mouse is then flipped over to the prone position for magnet placement.

#### Magnet placement

To a sterilized and shaved scalp, a 1-1.5cm midline incision was made using sterile scissors and forceps. On the stereotaxic set up, the mouse head was rotated 40-45 degrees to reveal the right temporal muscle. Using fine scissors and forceps, the margin where the temporal muscle attaches to the skull bone was cut and the flap of temporal muscle carefully retracted to expose the temporal bone below. Bleeding was controlled using sterile cotton swabs and frequent flushes of saline. As the supraorbital vein and retro-orbital sinus sit right along this margin, surgeons were mindful of extensive resection to avoid excessive bleeding. The branches of the MCA were visible, climbing upward from behind the zygomatic arch, below the temporal bone. A 2mm area was thinned over the main trunk and distal branches of the MCA using a dental drill equipped with a ¼ mm drill burr on medium to low speed. Alternating every few seconds with saline-soaked cotton swabs to remove bone powder and to prevent overheating of the skull by drilling, the skull over the trunk of the MCA was thinned as close to the zygomatic arch as possible, using rapid strokes and light pressure while holding the drill-bit nearly parallel to the bone. The skull is sufficiently thinned to 70µm once the vessel is clearly visible without the addition of saline. The attractive magnet (1.59 x 0.79mm thick N52 NdFeB axially magnetized Ni coated magnet; Zigmyster magnets Cat No. D116-132N52) was placed flat over the thinned region and glued down using Loctite gel superglue. Ideally the magnet was placed on the ascending trunk of the MCA and any probe measuring cerebral blood flow was placed on downstream branches or just adjacent to the magnet on the trunk of the vessel but further downstream if no branches were visible. Note that for reperfusion of vessel, the magnet and glue could be easily lifted by forceps.

#### Particle delivery and MCA occlusion

While the animal is still in the prone position, and immediately before injection of particles, the CCA was temporarily ligated by gently pulling on the implanted nylon string and taping the ends to the stereotaxic setup, so both strings were taut. CBF reduction of ∼20% is expected here. Thrombin-BSA conjugated particles suspended in physiological saline were sonicated for 10 minutes prior to delivery. Warm water was used to dilate the mouse tail vein, followed by an ethanol surgical wipe to the tail immediately prior to injection. Using a 1ml syringe and 30G needle, 250 µl (2.5 mg particles/mouse) sonicated particles were injected by tail vein. To monitor effective drop in blood flow, Doppler flowmetry (Perimed) or Laser Speckle Flowmetry (Omegazone) was used to record a sustained CBF decrease of 85% or greater for 30 minutes.

After 30 minutes, the CCA was released from ligation and magnet was removed gently by forceps. The ligation was loosened by cutting the two tense, taped silk sutures. If continuously monitoring cerebral blood flow, one can carefully slide out the remaining relaxed string out from under the carotid still in the prone position, slowly using forceps. If not measuring blood flow, animal can be flipped to the supine position to slide out remaining suture from behind carotid. Animal was then closed for recovery using 6.0 USP nylon suture string on the neck and sterile wound clamps on the scalp. Lidocaine topical analgesic was applied to surgical sites. A prepared, clean warm recovery cage is equipped with wet food and gauze for recovery. Animals are placed in physiologic recovery chamber (Darwin chambers, 31°C) for a minimum of 48 hours and up to 7 days until ready to image or be sacrificed. Following surgery, Buprenorphine at 0.5 mg/kg is administered every 6-8 hours subcutaneously for the first 72 hours following stroke.

Sham animals underwent the identical surgery to the tMP stroke groups, including skull thinning and transient CCA ligation, without placement of the attractive magnet. Sham animals were injected with either magnetic thrombin nanoparticles (tMP) or physiological saline as indicated.

#### Note

Investigator may opt to use rodent ear-bars to stabilize the animal’s head during surgery, however it is not absolutely necessary for this procedure and strict use of the stereotax and ear-bars may impede in surgical flexibility as the head will be rotated by about 40 degrees, making proper placement of the bars a challenge. Adding hand stabilization techniques, however most comfortable for the surgeon, was found to be most beneficial in avoiding error during surgery, particularly with thinning the temporal bone and magnet placement.

### Awake stroke procedure

Prior to surgery, mice were acclimated to a tail vein restrainer by one week of handling by researchers once daily for 5 minutes each. Handling is defined by holding mice, stroking fur and tail and allowing animals to climb on hands of researchers. Handling is followed by 4 days of placing animals individually into the tail vein restrainer for one minute.

#### Unilateral CCA stenosis by coil

Animals were anesthetized (Isoflurane induction 4-5%, maintained at 1.5%). Transdermal Bupivacaine 0.25% solution is administered to neck and scalp prior to incision. With animal supine and using sterile surgical technique, a 1cm incision was done on shaved and cleaned skin over trachea, without cutting trachea or muscles. Using sterile cotton swabs with saline, neck muscles were pulled apart to reveal trachea and surrounding vessels. Right CCA was isolated and a 10cm long 4-0 US silk surgical suture was fed below the rostral segment of carotid. A 0.18mm coil (Sawane Spring, Cat.No. SWP-A 0.18) was placed around the carotid, using the surgical suture to maneuver the vessel around the coil. Once the coil was properly placed around the CCA, the surgical suture was pulled out from below the carotid. Cerebral blood flow measurement using LSI over the surface of the skull and comparing to the contralateral hemisphere was done to confirm a 20-30% drop in CBF as a result of the carotid stenosis. Once confirmed, the neck skin is re-sutured with the coil left in place and lidocaine applied. Flipped to the prone position, the magnet was glued over a thinned temporal bone in the same technique outlined in the tMP stroke procedure. Once the magnet is well-glued, the scalp was closed using wound clips. Lidocaine was applied over incisions. Once fully ambulatory and exhibiting normal feeding and grooming behavior (by 1-2 hours) mice were prepared for tMP injection as described with the exception that mice are injected by tail vein while awake. Prior to tail vein injection, 0.25% bupivacaine was injected at the base of the tail and topical lidocaine was applied over the length of the tail. Once in the restrainer, the mouse tail was carefully warmed (e.g. with a heat lamp or warm water) to cause vasodilation, increasing ease of vascular access. Once veins were clearly visible and dilated, 250 µl (10 mg/ml) of sonicated magnetic BSA-thrombin particles in saline was injected using a 1ml syringe and 30G needle. Following particle injection, animals were immediately placed in a warmed cage and soon after transferred to a designated regulated physiologic recovery chamber. Following surgery, Buprenorphine at 0.5 mg/kg was administered every 6-8 hours subcutaneously for the first 72 hours following stroke.

### Circadian experiments

Mice underwent the awake stroke procedure at one of two dedicated times in a 12-hour period. Mice had surgery mid-morning or late afternoon, followed by tMP injection 1-2 hours later. Dedicated stroke (tMP injection) times were at either Zeitgeber time (ZT) 6-7 during their sleep inactive cycle (mid-day) or ZT 13-14 during their wake active cycle (early evening). Mice were injected as a group following recovery of surgery. “Anesthesia” mice underwent the same surgery outlined for the awake stroke procedure of coil and magnet placement and allowed to recover but instead underwent anesthesia a second time for tMP injection for 10 minutes during the same dedicated Zeitgeber times. As in previous, mice were immediately placed in a warmed cage and transferred to a regulated recovery chamber and following surgery Buprenorphine was administered at a dose of 0.5 mg/kg every 6-8 hours. Mice were sacrificed 48 hours later and assessed for stroke volume using Nissl staining.

### Assessing systemic distribution of tMP by Magnetic Resonance Imaging

Magnetic resonance imaging (MRI) was performed on a BioSpec 70/30 USR 7.0 Tesla Small Animal MRI at the Citigroup Biomedical Imaging Center (Weill Cornell Medicine) with a 200 mT/m gradient amplitude, a 640 mT/m/s slew rate and a 20 cm inner bore diameter. Whole body T1 isotropic, T2 weighted and T2Star mapping was done in animals injected with saline, tMP, and 30 minutes following tMPS. Animals were anaesthetized with 1.5-2% isoflurane during imaging. Raw DICOM data was processed using Fiji Image J software.

### Assessment of physiological tolerance of magnetic particles *in vivo*

Femoral artery catheterization was performed to collect arterial blood for pCO_2_, pO_2_ and pH measurement in tMPS and tMP sham injected animals before and following injection (Siemens Epoc Blood Analyzer System) (N=5/group). Animals were under 1.5% isoflurane anaesthesia. Arterial pressure was recorded by femoral artery catheterization using a pressure transducer (ADInstruments PowerLab data acquisition system and ADInstruments LabChart software for analysis). Given, the T2 weighted MRI images revealed accumulation of particles to the liver, liver serum chemistry was done at 4h, 24h, 48h and 7 days for alkaline phosphatase, alanine transaminase, aspartate transaminase, total serum protein, triglycerides, and total bilirubin by collecting blood from cheek puncture. Serum chemistries were performed by the Center of Comparative Medicine and Pathology at Memorial Sloan Kettering Cancer Center.^65^ Serum levels of MCP-1, MIP1α, IL-4, IL-10, IL-6, IL-1β, IP-10, RANTES, and TNFα were measured by an U-PLEX multiplex assay (MSD custom order) and read by a MesoScale plate reader. Serum was collected from stroke, particle sham and saline groups at 4 h, 24 h, 48 h and 7days after intervention.

### Cerebral blood flow measurement

Day of surgery cerebral blood flow was measured using Laser Doppler Flowmetry (LDF). Downstream flow from the distal branches of the MCA was measured using an optical fiber paired with the Perimed Master Probe 418. The fiber/probe was placed with the flat tip of the fiber flush on the temporal bone over the vessel downstream of the magnet. Both the glued fiber/attached probe and magnet remained glued on the temporal bone for the entirety of blood flow recording. Pairing (Loctite) gel glue and (Insta-set) glue accelerator was necessary to facilitate precise placement. Laser Doppler was continuously measured to give recordings at baseline, during the tMP injection/occlusion and magnet removal/reperfusion. Raw data files were normalized to a baseline value of 100 and every 5 seconds of linear recording was averaged using a custom R script. Data was plotted for corresponding the 5 minutes baseline recording, 10 minutes carotid ligation, 30 minutes occlusion and 30 minutes reperfusion performed in surgery. Linear recordings were binned to reflect an average for each time segment for each animal and plotted as bar graphs. Note that occlusion and reperfusion groups were binned using the last 10 minutes of recordings and this segment was used for statistics.

Laser Speckle Imaging (LSI) was also used to measure whole brain spatial and temporal dynamics of cerebral blood flow to both hemispheres during tMP stroke and in the days following to model reperfusion to the MCA after the magnet is removed.^66^ These chronic imaging experiments were performed on an Omegazone OZ-2 Laser Speckle Tissue Blood Flow Imager with laser unit and CCD camera. Anesthetized animals (1.5-2% Isoflurane) were prepared for tMP stroke as described previously including the nylon suture placed behind the CCA and magnet placement, prior to preparing the skull for LSI.

Prior to imaging, whole skull thinning was performed with the dental drill. With animal securely attached to ear bars in the stereotaxic table and scalp still open from previous magnet placement, scalp and white periosteum was trimmed laterally to expose the entire dorsal surface of the skull. Scalp skin was glued to edges using Vetbond. Note that the temporal muscle was already removed for the placement of the magnet and the magnet will still need to be removed for MCA reperfusion. This side of the scalp was left partially unglued until procedure was finished and tissue glue on this side was added when animal was ready to move to recovery. Thinning of the skull to approximately 90 µm thickness on both hemispheres was done to resolve the MCA for imaging. Daily repeated thinning was needed to control for bone re-growth as animals were imaged for several days. Prior to imaging, bacteriostatic light scattering gel was applied using a cotton swab. The gel is removed using a cotton swab and a sterile saline rinse is applied to clean the surface of the skull before the animal can recover. LSI was done longitudinally providing recordings at baseline, during the CCA ligation, magnetic particle injection/occlusion, and magnet removal/reperfusion. Animals were re-anesthetized and imaged up to five days following surgery. For each time point, LSI images were obtained every 1 second for duration of the procedure. Frames with significant artifacts obscuring the MCA were deleted and LSI images were averaged over 1 minute per each time point and using Fiji ImageJ software a region of interest was drawn around the major MCA branch on each corresponding cortex and compared with the identical region on the contralateral side measuring changes in pixel intensity.

### Nissl staining for infarct volume measurement

Brains were prepared fresh and flash-frozen in an isopentane bath. Cryostat (Leica CM1850) slices of coronal sections at 30 µm thickness were collected every 250 µm. Sections were fixed in 4% paraformaldehyde in 0.1M phosphate buffer for 10 minutes. Following fixation, sections were rinsed in 0.1M phosphate buffer for 5 minutes and then incubated in Cresyl Violet solution for 12-15 minutes. Sections were then placed in water for 2 minutes before dehydration in sequential baths of 50%, 75%, 95% and 100% ethanol, 10 seconds each. Sections were cleared in two sequential 2-minute incubations of Xylene before mounting with DPX permanent mounting solution. Infarct volume determination was done using ImageJ software. Using the freehand selections tool in ImageJ, the perimeter of infarct was traced and area measured and converted to mm^3^. The contralateral hemisphere was used to correct for edema as previously described.^12^

### Immunohistochemistry

At 2, 7, and 14 days following stroke, animals were deeply anesthetized using sodium pentobarbital (150mg/kg) and transcardially perfused with ice cold Phosphate Buffered Saline (PBS) and 2% Paraformaldehyde (PFA) fixative. Isolated brains were kept at 4 °C in the 2% PFA overnight before moving to 30% Sucrose for two days. 18 µm sections were cut by cryostat (Leica CM1850). Sections were rehydrated with 1X PBS and permeabilized using 0.5% PBS-T (Triton X100) for thirty minutes. Next, sections were blocked for 2 hours in 5% normal donkey serum in 0.1% PBS-T. Sections were incubated in 0.1% PBS-T, 1% normal donkey serum and with corresponding primary antibodies for two days at 4 °C. Primary antibodies included anti-MAP2 (1:200, rabbit polyclonal, abcam Cat. No ab32454), anti-Iba1 (1:500, rabbit polyclonal, Wako Sigma Aldrich Cat No. SAB5701363), anti-GFAP (1:200, rabbit polyclonal, abcam Cat. No ab7260). Primary antibodies were removed by washing sections twice with 1X PBS for 5 minutes. Sections were incubated for two hours at room temperature shielded from light in 1% normal donkey serum, 0.5% PBS-T and corresponding secondary antibodies in FITC goat anti-rabbit (abcam Cat. No ab6717), Cy5 goat anti-rabbit (Thermofisher Cat. No A10523) or Cy3 goat anti-rabbit (Thermofisher Cat. No A10520) at 1:100. Sections were washed twice with 1X PBS for 5 minutes. Sections were mounted using Fluorosave containing DAPI (Sigma-Aldrich Cat No. F6057).

Immunostaining using anti-fibrinogen antibody (rabbit anti-mouse polyclonal abcam Cat. No ab34269) and antibody for platelet marker CD41 (rat anti-mouse Biolegend Cat. No 133903) of the distal branches of the MCA and embedded clot were done on a vibratome-cut 40 µm free-floating coronal sections. Brain sections were selected from 24-well storage container and rinsed with 0.1M phosphate buffer to remove agar from vibratome. Sections were washed with 1X PBS twice for 5 minutes and then permeabilized with 0.5% PBS-T for 1 hour at room temperature. Sections were washed with 1X PBS twice for 5 minutes. Sections were then blocked with 3% normal donkey serum in 0.5% PBS-T for 1 hour at room temperature. Sections were washed twice with 1X PBS for 5 minutes. Sections were left in primary antibodies 1:500 anti-fibrinogen and 1:500 anti-CD41 overnight in 0.5% PBS-T at 4°C. Sections were washed with 1X PBS the following day. Secondary antibodies, Cy3 goat anti-rabbit (Thermofisher Cat. No A10520) and FITC goat anti-rat (ThermoFisher Cat No. 31629), were incubated at 1:100 for 2 hours in 0.5% PBS-T at room temperature, shielded from light. Floating sections were washed thoroughly with 1X PBS and mounted on slides. Slides and sections were allowed to dry before mounting coverslips using FluoroSave (Sigma-Aldrich F6057).

### Fluorescent Activated Cell Sorting

Mice were anesthetized with sodium pentobarbital (150mg/kg) intraperitonial. Single cell suspensions of mouse cerebral cortex were prepared after transcardial cold PBS (containing 2U/ml Heparin) perfusion. The cerebral cortex was dissected and placed into tubes containing 2ml digestion solution (HBSS, 10mM Hepes pH 7.4, 1mM MgCl_2_, 50U/ml DNAse I) on ice. Once all brains were collected, 50µl of Dispase (Worthington Cat No. 2104, 32U/ml) and 50µl Liberase DH (Roche Cat No. 12352200, 2.5mg/ml) was added to each sample. Tissues were homogenized using a tissue dissociator (gentleMACS, m_brain_01 program, Miltenyi Biotech). Samples were then placed in an orbital shaker for 45 minutes at 37°C at 100 rpm. Tissues were again homogenized using a tissue homogenizer (gentleMACS, brain_03 program) and transferred to 15 ml falcon tubes on ice. Samples were centrifuged at 1000*g* for ten minutes at 4°C. Pellets were resuspended 1ml in 30% Percoll (Cytiva Cat. No 17089101) in HBSS/HEPES and resuspended 25 times using a p1000 pipette. 9ml of 30% Percoll was then added to each sample, mixed by inversion, and samples were spun at 1750 rpm for 15 minutes at 4°C. Myelin was removed by aspiration using a Pasteur pipette. Supernatant was removed and pellets were individually resuspended in 10ml HBSS/HEPES. Samples were spun at 1000*g* for seven minutes at 4°C. Supernatant was discarded and pellets were each resuspended in 1ml FACS (2% Fetal Bovine Serum, 0.05% NaN_3_ in 1x PBS) buffer into FACS tubes. 3ml of FACS buffer was then added to each sample and samples were spun at 1500 rpm for 6 minutes at 4°C. Supernatant was discarded and samples were blocked in 50 µl FACS buffer containing CD16/CD32 (Biolegend Cat No. 101301, Clone 93, 5 ng/µl) for 10 minutes. Samples were then stained with 5 µl of corresponding antibody mix. The following antibodies and channels were used to differentiate leukocyte populations CD4 BV510 (Biolegend Cat. No 100553, Clone RM4-5 0.038ng/µl), CD45 APC (Biolegend Cat. No 103111, Clone 30-F11, 1.36 ng/µl), Ly6C BV711 (Miltenyi Cat No 130-099-353, Clone 1G7.G10, 0.39 ng/µl), TCRγδ PE (Biolegend Cat. No 118124, Clone GL3, 0.39 ng/µl), CD8a AF700 (Biolegend Cat. No 100729, Clone 53-6.7, 0.03 ng/µl), NK1.1 PE-Dazzle (Biolegend Cat No. 156517, Clone s17016D, 1.61 ng/µl), CD19 PE-Cy5.5(Invitrogen Cat No. MHCD1918, Clone Sj25-C1, 0.9 ng/µl), TCRβ PE-Cy7 (Biolegend Cat No. 109222, Clone H57-597, 0.39 ng/µl), Ly6G FITC (Biolegend Cat No. 127605, Clone 1A8, 0.61 ng/µl), CD11b APC/Cy7 (Biolegend Cat No. 101226, Clone M1/70, 0.08 ng/µl). Samples were stained for 15 minutes on ice and washed in 3 ml of FACS buffer. Samples were spun for a final time at 1500 rpm for 7 minutes at 4°C and then resuspended in 200µl of FACS for sorting. Samples were acquired on a NovoCyte Quanteon flow cytometer and analyzed using FlowJo version 10.7 software. Discrimination gates were established using isotype controls and fluorescence minus one experiments.

### Neurological deficits

Behavior tests were performed to measure the functional neurological deficits of mice following tMP stroke. Animals underwent hanging wire and corner test recordings.^23^ Two days following injury, mice are suspended on an outstretched, taut wire between two poles roughly 60 centimeters above the ground or table. Mice must initially grab the wire with both fore-paws and allowed to “hang” and are given 60 seconds to reach either of the two poles. The “latency to fall” is the amount of time the animal can remain suspended without falling. After 1 minute, the trial is over. Animals reaching either pole are automatically given a score of 60 seconds for the trial. Three trials are performed for each mouse with 10 minute intervals. The corner test was used to establish sensorimotor deficits and motor asymmetries following injury. The testing setup included two darkened walls angled at 30 degrees to a corner. At the junction of the two walls, a small amount of light should be allowed to pass through, and food pellets can be placed behind the walls to motivate the animal to enter the corner. Animals are placed between the two walls and allowed to approach the corner and then rear upwards, both palms on a wall, and turn back around. Investigators record the direction of rearing/turning. Trials in which the animal does not rear upward while turning are discarded. Each animal performs consecutive 10 trials. Animals also performed the adhesive removal test at 3 days following stroke for assessment of sensorimotor deficits^26^. Animals receive two adhesive tape strips to their forepaws and timed for 120 seconds in a clear testing box equipped with a camera. Contact time is defined as the point in which the mouse recognizes the presence of the tape strips (ex. shaking paws or touching tape to the mouth). Removal of the first and second pieces or tape are recorded for the moment the tape touches the bottom of the clear container. Animals were trained three days before stroke to establish a baseline reading and re-tested at 3 days following stroke.

### Adaptation of tMP Stroke for Multiphoton Imaging

All steps of the surgical preparation for magnetic particle stroke were done prior to preparing the skull with a thin-skull window for imaging. The original scalp incision which was done for magnet placement was elongated as needed to expose the entire occipital bones. Using forceps and surgical scissors, the periosteum was trimmed laterally to the edges of the temporal bone. Scalp skin was glued to the edge of where the temporal muscle inserts onto the temporal bone and the tissue on the posterior of the neck using Vetbond tissue glue, exposing whole dorsal skull surface. Note that one will have already removed parts of the temporal muscle for placement of the magnet and may choose to remove the magnet after the 30-minute occlusion. Therefore, this portion of the scalp remains open until imaging procedure is finished and the animal is ready to move to recovery. Vetbond can be used to close the scalp. Mice are secured in the stereotaxic setup prior to thinning and placing the headplate using rodent ear bars. Using a ½-mm burr on medium to low speed, the parietal bone was lightly thinned on the ipsilateral side of the occlusion to slightly flatten the area where the headplate will be added. The skull was swabbed every few seconds with a saline-soaked cotton swab to avoid overheating and remove bone fragments. The distal branches of the MCA are visible in C57Bl/6J mice ascending below the surface of the temporo-parietal bone, roughly perpendicular to where the magnet was placed over the MCA on the temporal bone. A 2-mm by 2-mm region was mapped over where the vessel is seen under the parietal bone. Investigators thinned through the cancellous layer of bone in this 2-mm by 2-mm region using low drill speed and feather-light strokes, always alternating every few seconds with saline and cotton swabs. At the point where the pial vessels are visible under moist saline-soaked bone, white spots in the spongey bone layer will show on the surface of the skull. The area was continuously thinned until these spots disappeared, indicating the inner plate of the parietal bone has been exposed. Extreme care is taken here to not apply unnecessary pressure and avoid overheating of the bone. At this point a small droplet of cyanoacrylate glue was added over the thinned area and a 2-mm round glass coverslip was placed flat over the glue. The glue and coverslip will prevent bone re-growth and imaging aberrations during chronic imaging studies. If imaging once, the thin-skull window can be left bare and submerged in water when imaging. After coverslip placement, a custom headplate was affixed with dental cement (C&B Metabond, Parkell) to the lateral aspect of the parietal bone just superior to the temporal bone.^67^ The headplate was positioned with the screw points facing laterally, roughly perpendicular to the temporal bone, leaving the temporal bone and glued-on magnet below clean of dental cement. Dental cement was built up in a circle around the coverslip to prevent bone re-growth, allowing cement to slightly dry and thicken before placing the titanium headplate.

### Two photon imaging of mice undergoing tMP stroke

Mice were kept anesthetized with 1.2-1.5% isoflurane in mixed oxygen and nitrogen to target a ∼1Hz respiratory rate while keeping the core body temperature at 37°C with a feed-back controlled heating pad. Mice were stabilized by attaching the headplate to a metal frame on an optical breadboard. 70kDa Texas Red-conjugated dextran (50 µl, 2.5% w/v dissolved in saline, Invitrogen) was injected retro-orbitally to visualize the vasculature just prior to imaging. Leukocytes and blood platelets were labeled with Rhodamine 6G (0.1 ml, retro-orbitally, 1 mg/ml in 0.9% saline, Acros Organics). Hoechst 33342 dye was also injected retro-orbitally (50 µl, 4.8 mg/ml in 0.9% saline, Thermo Fisher Scientific) to label leukocytes and distinguish them from platelets. Before imaging, the surface of the coverslip was gently cleaned with a moist cotton swab. Red blood cell speed and vessel diameter changes were monitored at baseline, following CCA ligation, during injection of tMP, and following removal of magnet and CCA ligation. These steps were done while the mouse was secured to the imaging setup. Animals’ neck tissue was re-sutured following imaging and the mouse was allowed to recover in ambient stroke recovery chamber for future imaging.

Imaging was performed on a two-photon microscope (FVMPE, Olympus). Excitation pulses came from a solid-state (InSightDS+ Spectraphysics) laser at 800nm. Fluorescence was collected through band-pass emission filters (Hoechst, 410-460nm; Rhodamine 6G, 495-540nm; Texas Red, 575-645nm) and delivered to photomultiplier tubes. All image stacks were collected using Fluoview software (Olympus), where each hyperstack movie consists of the 3 fluorescent layers. To visualize brain architecture and identify vessels of interest, a 300 µm deep wide field map was taken using a 5X objective (0.28 NA, Olympus). Once vessels of interest had been identified, we used a water-submerged 25X objective (SLPlan N 1.05 NA, Olympus) to perform all line scans and acquire z-stacks. Red blood cell velocity was measured by repetitive unidirectional scans across the central axis of the vessel at 0.85 kHz for 60 seconds. Red blood cells were seen as diagonal dark streaks. The slope of these individual streaks is inversely proportional to blood cell speed. Scans were analyzed using MATLAB code with a Radon transform-based algorithm computing line integrals of the g(x,y) 2D image at various angles.^68^ All remaining images and movies were processed using ImageJ software.

### Statistical analysis

Data was analyzed for normal distribution using the Shapiro-Wilk normality test. Group-wise comparisons were performed with unpaired t-test, one- and two-way ANOVA, or Kruskal-Wallis test as appropriate. Statistical analysis was performed using Prism 10 (GraphPad Software).

## Acknowledgments

We thank all members of the Anrather and Iadecola laboratories for helpful discussion. This work was supported by NIH grants R01NS081179 (J.A), R01NS132493 (J.A), and the Leducq Foundation (StrokeIMPaCT Network; J.A). The generous support of the Feil Family Foundation is gratefully acknowledged.

## Author Contributions

K.M and J.A conceived the study with the input of C.I. K.M performed the experiments and analyzed the data with contributions from S.A, A.A, and N.R. S.A performed two-photon imaging and contributed to imaging analysis. A.A performed blood gas measurements. N.R performed some stroke experiments. K.M and J.A wrote the original draft. C.I revised the manuscript. All authors read and approved the final manuscript.

## Declaration of interests

C.I serves on the scientific advisory board of Broadview Ventures. The other authors declare no competing interests.

## Notes

### Summary of Updates

Figure 3 imported improperly. Revision has correct figure.

