## Supplementary figures and images for "A minimally invasive thrombotic stroke model to study circadian rhythm in awake mice"

### Supplementary figures

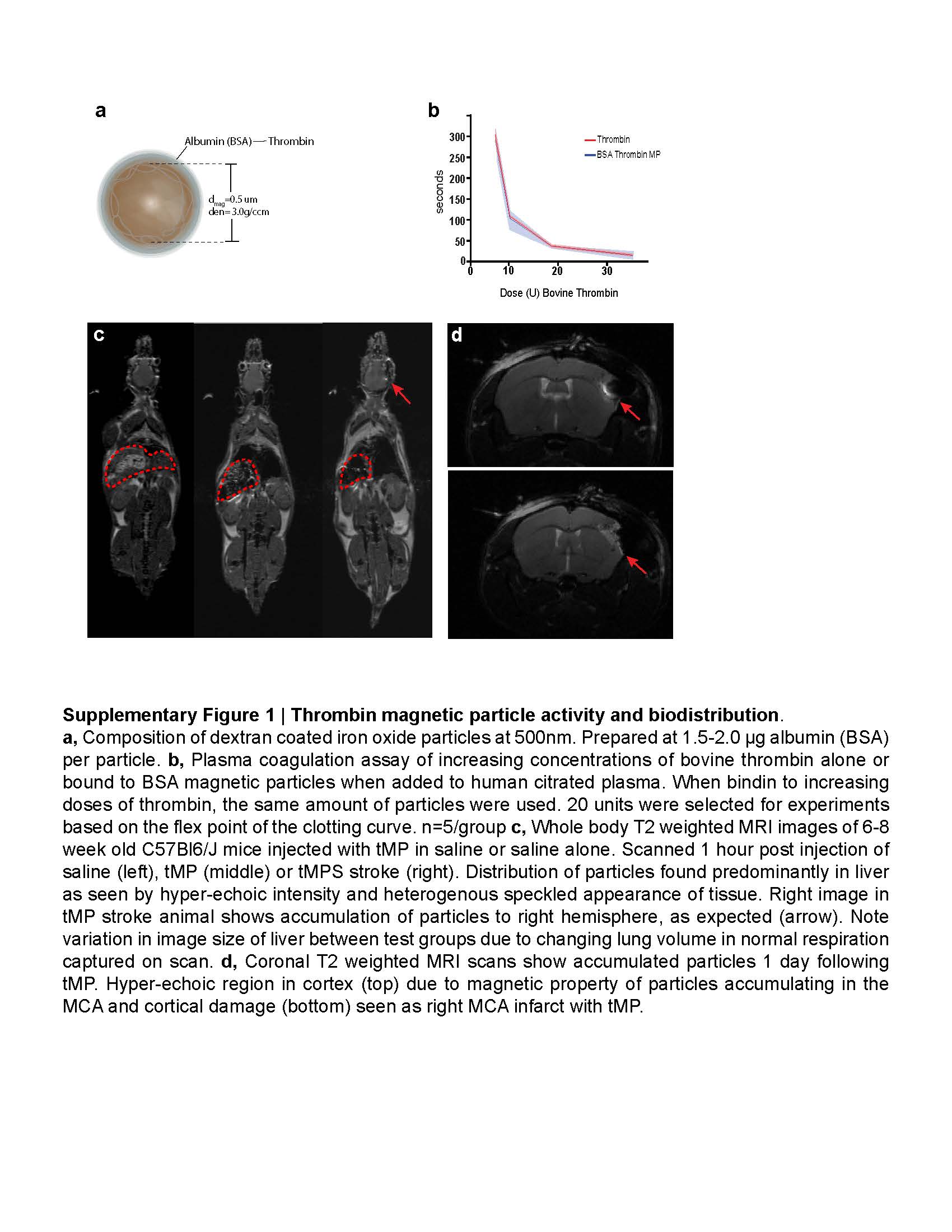

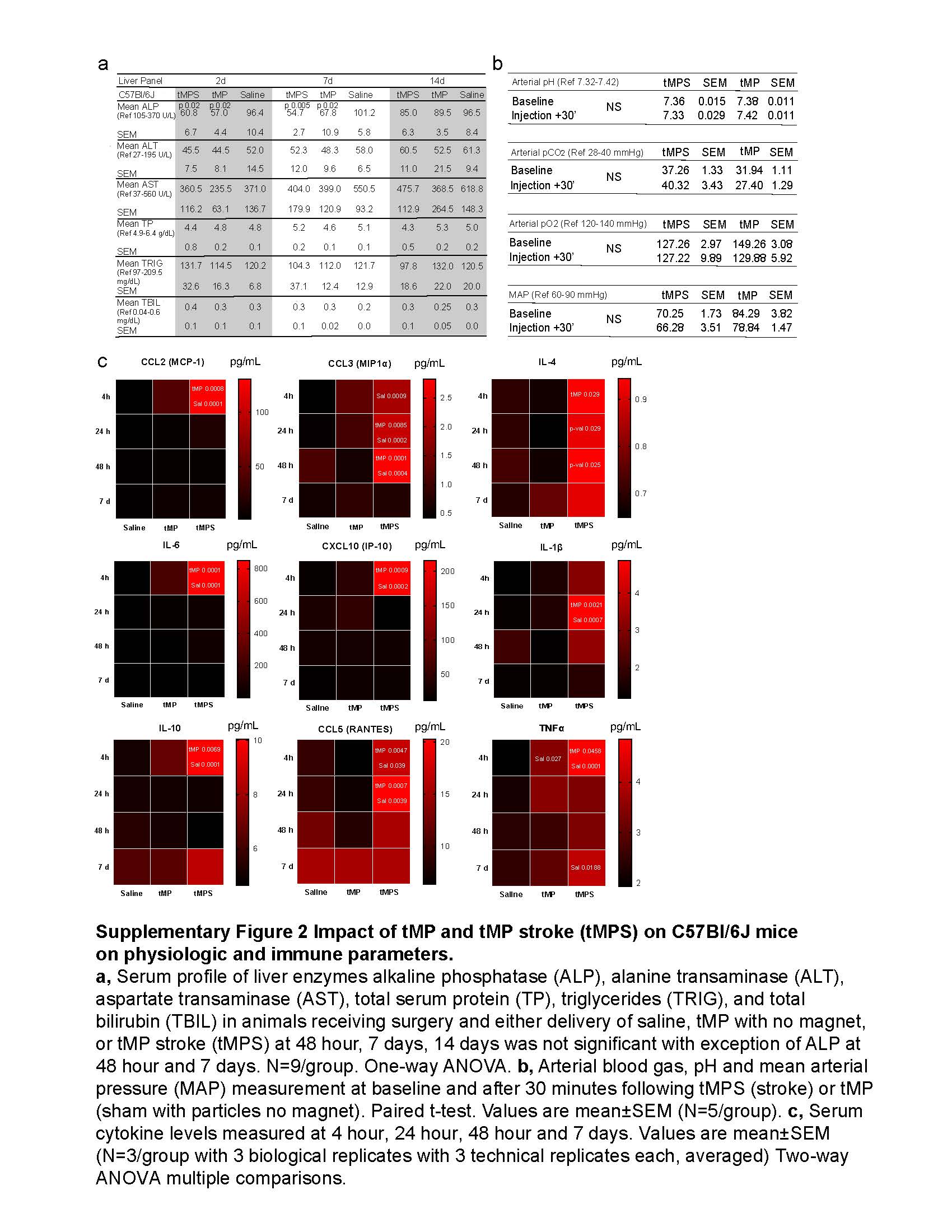

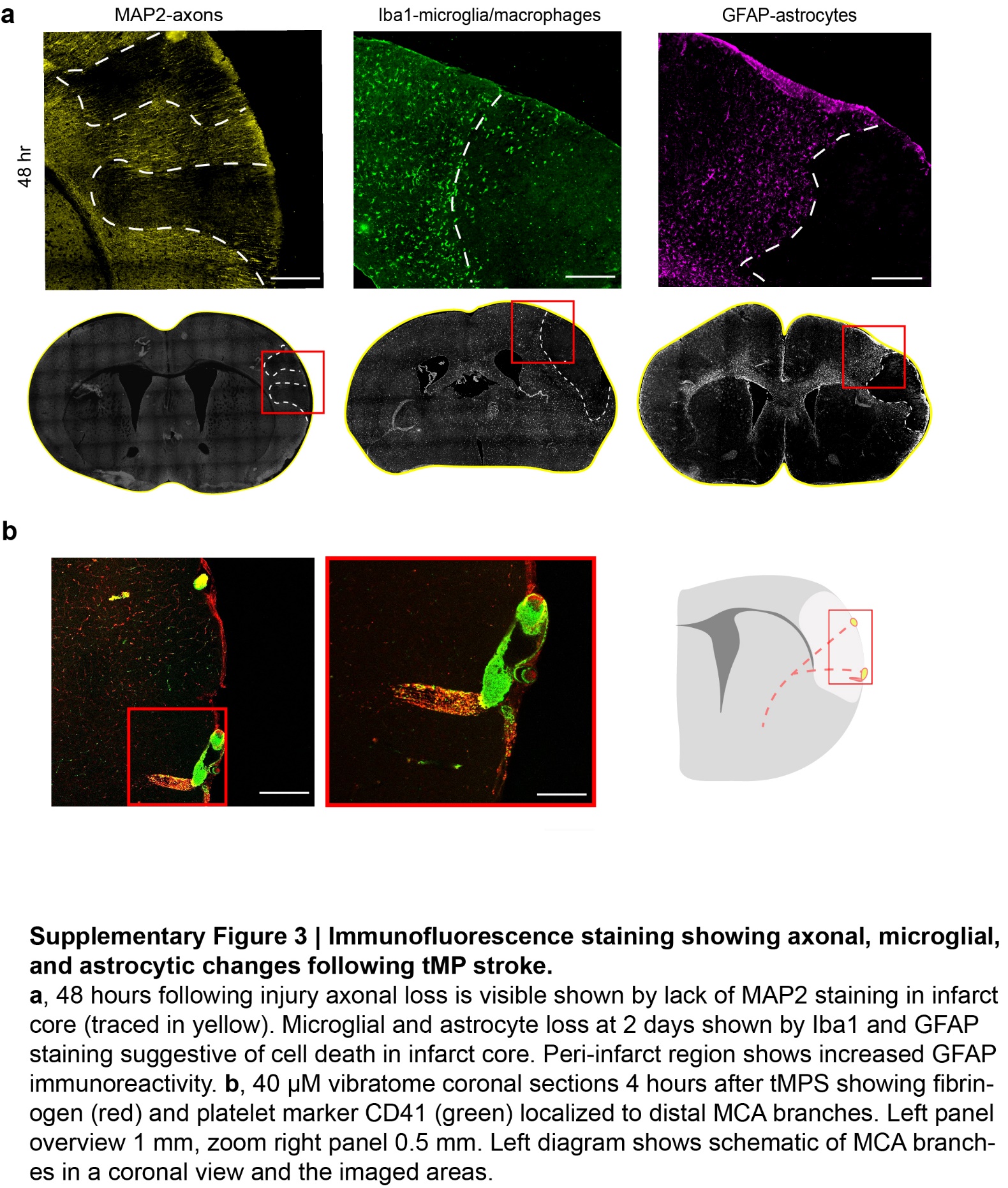

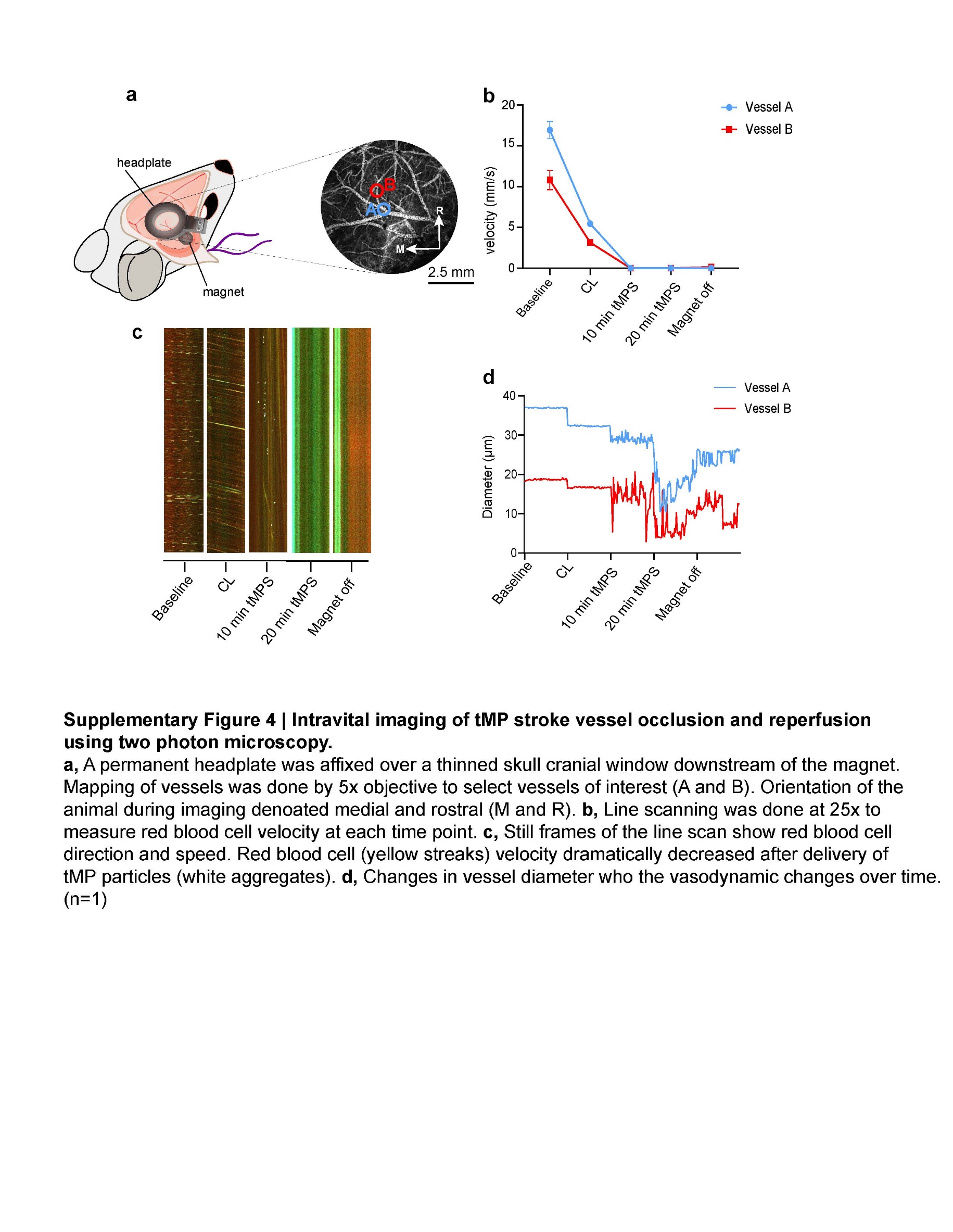

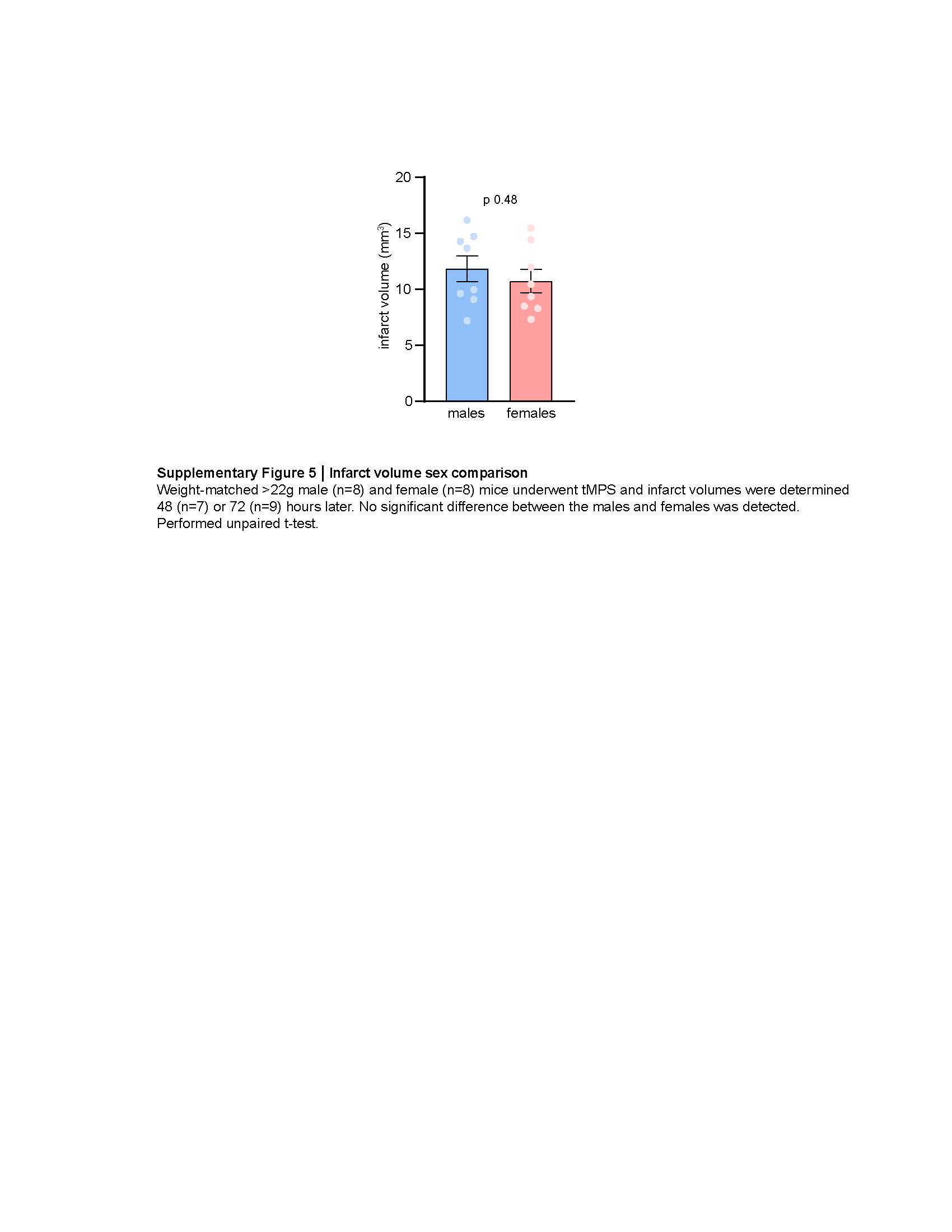

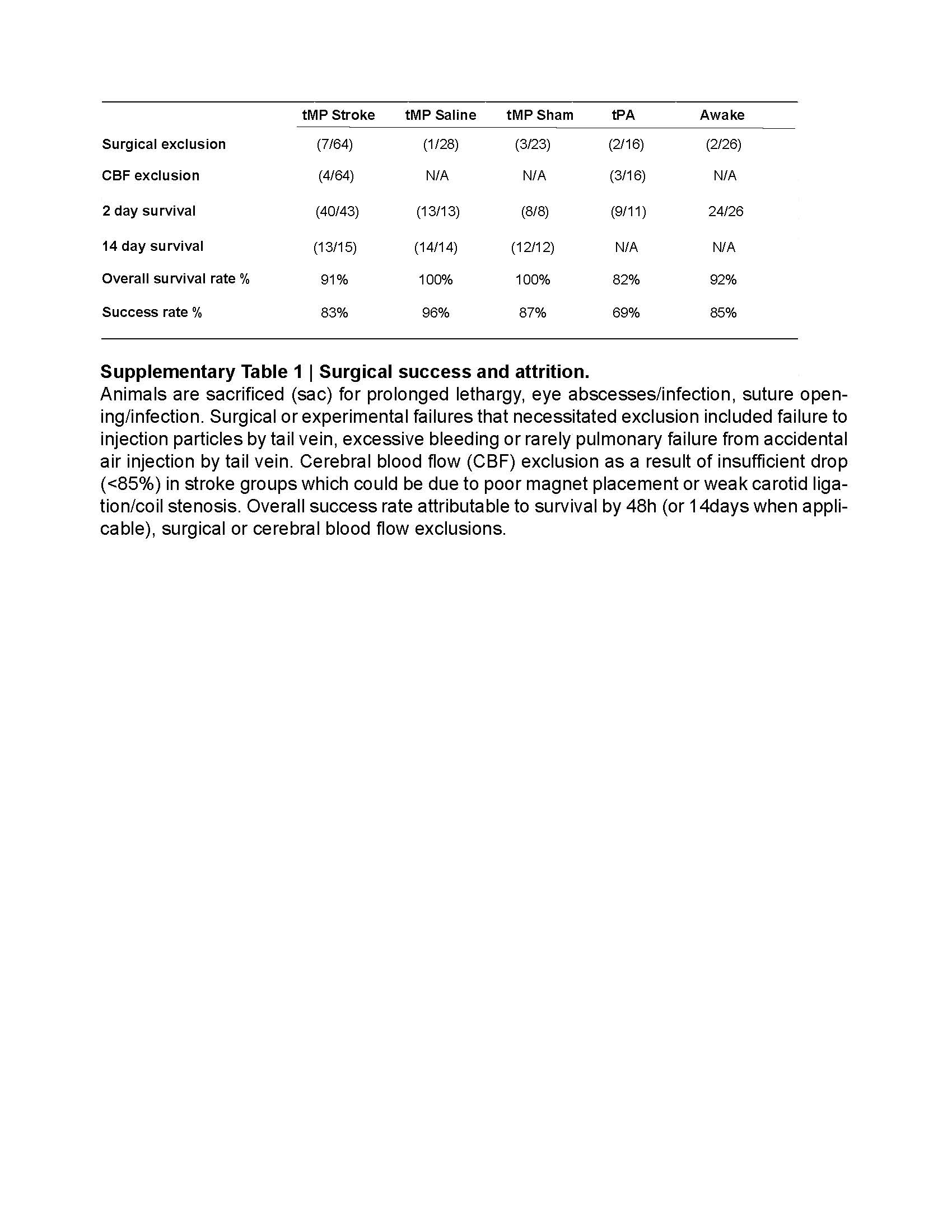

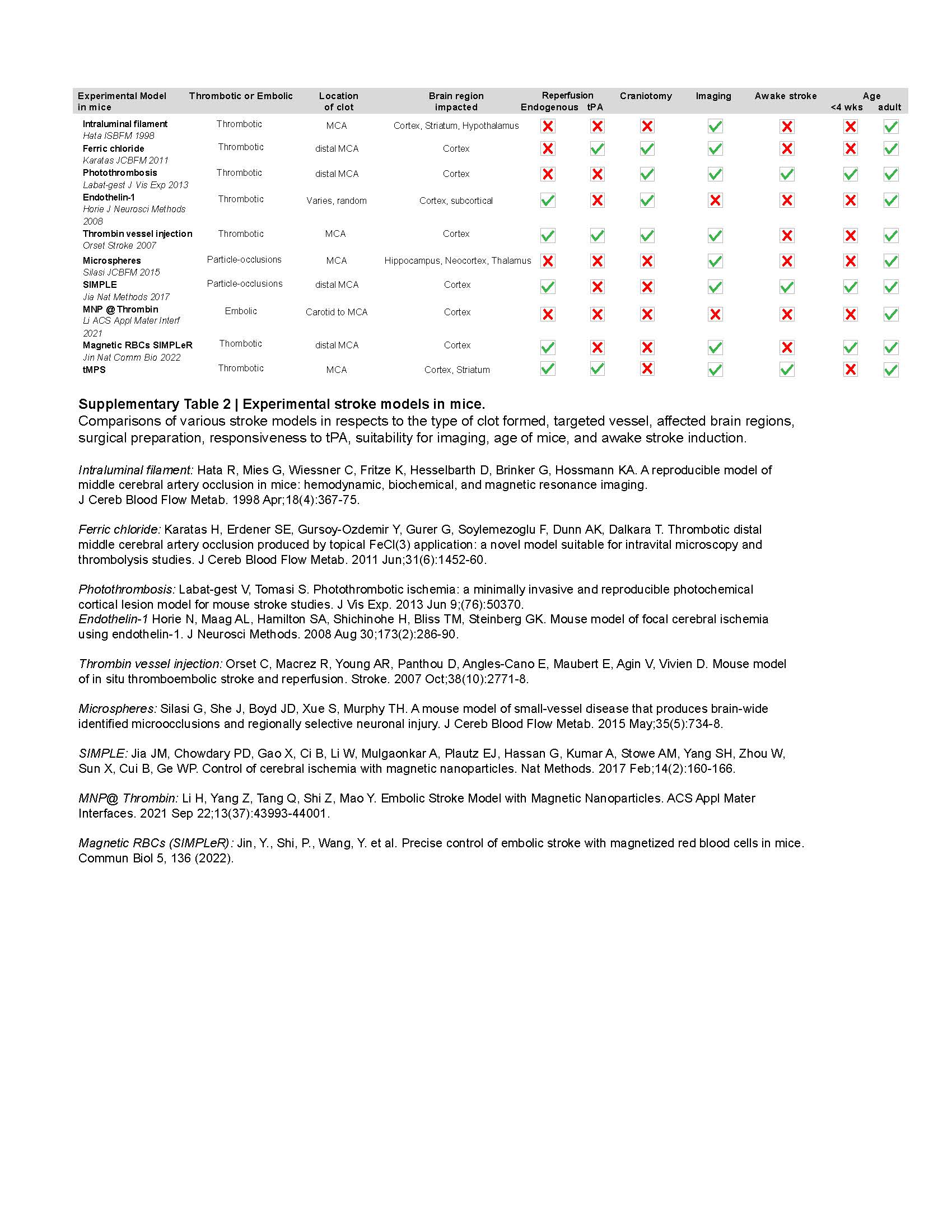
